# Revealing the transcriptomic organization of extrasomatic interfaces using SCOUBI

**DOI:** 10.64898/2026.11.10.765731

**Authors:** Bhavay Aggarwal, Saumya Jain, Saurabh Sinha

## Abstract

Spatial transcriptomics (ST) generates substantial transcript signal outside segmented cell bodies, which is generally discarded by analysis tools. This is especially problematic for brain ST data, as neuronal mRNAs can be localized far from somata – in axons, dendrites, and synaptic compartments – and up to half of the transcripts in ST data may lie outside cell bodies. Here, we present SCOUBI, a computational framework that identifies axon-dendrite interfaces from extrasomatic RNA in ST data, revealing a hitherto overlooked layer of molecular function in neural tissues. It first partitions extrasomatic space into axon- and dendrite-enriched compartments based on known marker genes, using a neural network trained to maximize marker specificity and inter-compartment communication signals. It then defines interfaces as loci where predicted axonal and dendritic compartments converge and exhibit ligand–receptor colocalization. Applied to mouse brain, SCOUBI creates maps of axonal and dendritic compartments and their interfaces that align with known spatial patterns of neurite and synaptic distribution. It identifies transcriptomic differences between interfaces and the surrounding extrasomatic space, revealing region-specific signatures of neuronal signaling and synaptic plasticity. By comparing interface transcript enrichment with local somatic expression, SCOUBI identifies genes that preferentially localize at sites of intercellular communication rather than simply reflecting nearby cell bodies. SCOUBI reveals interface-associated transcripts beyond established synaptic genes, with molecular signatures reproducible across independent samples and imaging platforms. In summary, SCOUBI transforms a substantial and often-discarded aspect of ST data into an interpretable readout of subcellular organization and neural signaling.

## Introduction

Sites of neuronal communication, such as synapses, are distributed along axonal and dendritic processes that can extend hundreds of microns from the cell body. The molecular machinery at these distal sites is partly specified locally, as mRNAs are actively transported to axons and dendrites, where they can be translated on demand in response to signals such as changes in abundance of calcium ions (Rangaraju et al. 2017; Das et al. 2021; Sun et al. 2021). The dysregulation of mRNA localization is implicated in a broad range of neurological disorders (Fernandopulle et al. 2021; Jishi et al. 2021). Mapping which transcripts localize to distal sites in healthy and disease-associated neurons remains a critical experimental challenge. Current approaches typically rely on isolating distal subcellular compartments, followed by transcriptomic profiling (Cajigas et al. 2012; Hafner et al. 2019; Kaulich et al. 2025). While direct, these assays are prone to biases introduced by the methods used for biochemical fractionation and are difficult to perform in intact tissue – they destroy the spatial organization needed to understand the functional implications of distal localization (van Oostrum et al. 2023; Matsubayashi and Takano 2025).

Spatial transcriptomics (ST) provides an alternative way to study transcript localization in intact tissue. A consistent feature of ST data is the substantial presence of transcripts outside segmented cell bodies (unassigned RNA, or uRNA) (Wang et al. 2020; Ament and Poulopoulos 2023; van Oostrum et al. 2023). Although some of this “extrasomatic” signal is attributable to technical noise or segmentation errors, accumulating evidence indicates that much of it is biologically meaningful. A recent study using MERFISH (Wang et al. 2020) demonstrated that axonal and dendritic mRNA markers spatially correlate with immunostained neuronal projections in cultured systems, establishing their transcript localization as a tractable proxy for subcellular compartment (neurite) location and identity. In this way, ST provides capabilities that direct synaptic profiling does not, including molecular resolution within intact tissue at scale and without the need for specialized fractionation.

The above observations suggest that extrasomatic transcript patterns in ST data may be used to identify spatial regions enriched in axonal or dendritic compartments as well as loci where these two types of neurites come into close proximity. A subset of the axon-dendrite convergence loci is expected to harbor signaling interactions with detectable transcriptomic footprints (e.g., *Gls*-*Gria1* representing glutamate → AMPA interaction), potentially making them molecularly distinguishable from the surrounding extrasomatic space. Thus, a properly designed search for markers of axonal and dendritic compartments as well as ligand-receptor pairs, in spatial proximity to each other, may reveal sites of neuronal communication and allow us to characterize underlying mechanisms and their regional and condition-specific variations (Zheng et al. 2025). (Note: we use “ligand–receptor pairs” to broadly include genes whose expression serves as a proxy for an underlying ligand–receptor interaction (Cang et al. 2023; Zhao et al. 2023; Kumar et al. 2026).) Building on this idea, we define “interfaces” as spatially localized regions in extrasomatic space where axonal and dendritic compartments converge and express ligand-receptor pairs mediating neuronal communication. Interfaces are a bioinformatic construct intended for deciphering molecular signatures and spatial distributions of communication; they do not correspond to individual synapses but instead represent micron-scale extrasomatic loci consisting of axonal and dendritic compartments and enriched for synaptic signaling programs.

Here, we develop and use a computational framework to identify neuronal interfaces from ST data and characterize their molecular signatures and spatial distribution patterns. The new framework, called SCOUBI (**S**ignaling**-**informed **C**haracterization **o**f **U**nresolved **B**iological **I**nterfaces), first annotates extrasomatic loci as axon- or dendrite-enriched using pre-specified marker genes. In this step, it addresses a fundamental informatic challenge: *how to weight different marker genes* (which may not be equally informative) *in designating axon- or dendrite-enriched compartments?* It does so via a dual objective optimization that demands that the designated compartments be enriched in their respective marker genes *and* neighboring axonal-dendritic compartments exhibit statistically significant communication patterns. The second step of SCOUBI utilizes the predicted axonal and dendritic compartments to pinpoint loci of axon-dendrite convergence and significant ligand–receptor colocalization, which are then predicted as interfaces.

Applying SCOUBI to mouse brain imaging-based ST (iST) data, we show that predicted axonal, dendritic, and interface loci recapitulate known features of neuroanatomical organization. Transcriptomic analysis of interfaces recovers molecular programs associated with neurotransmission and structural remodeling, and genes enriched at interfaces relative to surrounding extrasomatic space are strongly linked to synaptic function. We also identify interface-associated transcripts beyond established synaptic genes, including those with regulatory and metabolic functions. We characterize how molecular signatures of interfaces vary across brain regions, finding that interfaces harbor both a pan-region conserved gene program and region-dependent transcriptomic diversity. By contrasting interface transcriptomes with those of nearby cell bodies, we identify transcripts that preferentially accumulate at putative sites of intercellular signaling beyond what local somatic expression would predict. These analyses reveal region-specific programs, including a dopamine-glutamate integration program at striatal interfaces and structural remodeling signatures in the hippocampus. Benchmarking against existing approaches (Salas et al. 2025; Yuan et al. 2025) for extrasomatic RNA analysis and systematic model ablations demonstrate that SCOUBI’s communication-informed design provides a more biologically informative view of the extrasomatic transcriptomic landscape. Together, these results, reproducible across different independent tissue samples and iST technologies, show that extrasomatic transcript distributions can reveal putative axon–dendrite convergence regions, as well as the spatial and molecular properties of neuronal interfaces.

## Results

### Overview of interface prediction framework

We first describe our SCOUBI framework for predicting extrasomatic loci enriched for axon– dendrite convergence and ligand–receptor colocalization. These loci, which we call “interfaces”, are inferred from cell-segmented Spatial Transcriptomics (ST) input, via two steps: classification of extrasomatic “bins” as axon- or dendrite-enriched, and interface identification.

The input data are first spatially binned at a resolution (configurable, typically a few square microns per bin) that strikes a balance between statistical power, reflected in transcript count per bin, and spatial granularity (**Figure 1a**). A single-layer feedforward neural network model is then trained to assign each bin a probability of axonal or dendritic enrichment based on marker gene expression (**Methods**). Model parameters encode the relative weights assigned to different marker genes. These are optimized for marker gene specificity (e.g., axonal bins are enriched in axonal markers, **Figure 1b**) as well as the extent of inferred signaling activity between neighboring bins of opposing assigned identities (**Figure 1c**), quantified using the communication z-score from CellWHISPER (Kumar et al. 2026) (**Methods**). The communication z-score is the statistical significance of the frequency with which a specific neuronal ligand– receptor pair is expressed in adjacent axonal and dendritic-enriched bins; the model seeks neurite identity assignments that reveal many significant pairs. The dual objective training allows the model to learn a weighted combination of marker features in an unsupervised manner, integrating marker gene signal with spatial communication patterns. This process of designating extrasomatic bins as axon- or dendrite-enriched is henceforth called “neurite classification”.

**Figure 1:**
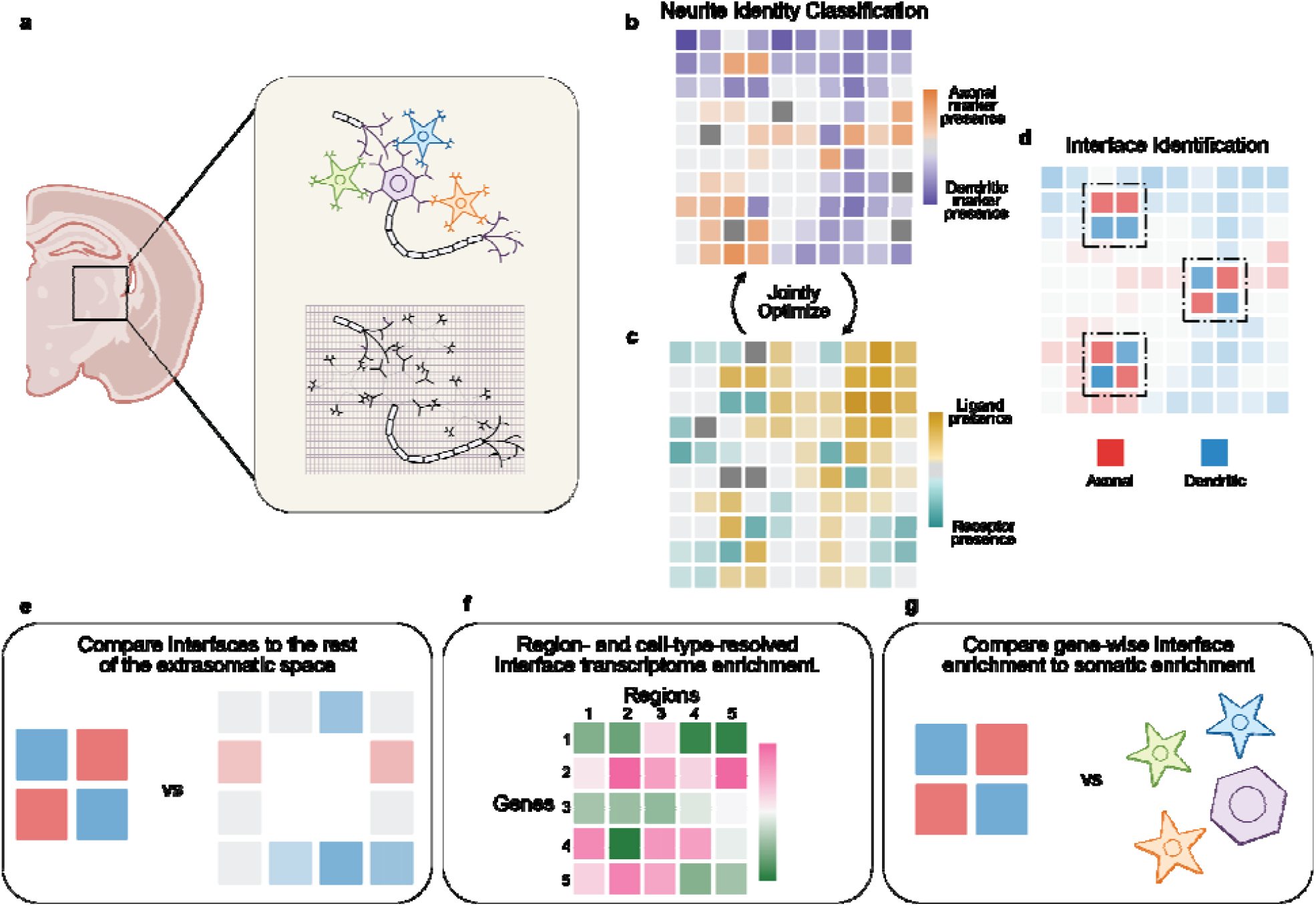
SCOUBI framework for neurite classification and interface identification. **(a)** ST data from a cell-segmented tissue section are spatially binned. Bins with majority transcripts assigned to a cell are excluded, leaving bins in the extrasomatic space as the substrate for analysis. Binning is at a resolution where each bin is as small as possible while containing sufficient transcripts for statistical analysis. **(b, c)** A feedforward model is jointly optimized for (b) marker gene specificity in axonal vs dendritic bins and (c) extent of inferred ligand-receptor communication between neighboring bins of opposing identity, quantified via the CellWHISPER communication z-score. **(d)** Interfaces are identified by scanning the annotated tissue with a grid, retaining regions containing two axonal (red) and two dendritic (blue) bins with sufficient ligand-receptor support. Multiple such interfaces are recovered across the tissue. **(e-g)** Downstream analyses enabled by SCOUBI include (e) comparison of interface transcriptomes against the broader extrasomatic space, (f) gene enrichment analysis for interfaces associated with specific regions and cell types, and (g) gene-level comparison of expression in interfaces versus somatic expression.

After neurite classification, interface regions are identified by scanning for loci enriched for axonal as well as dendritic bins using a 2 ×2 grid (**Figure 1d**). Such a grid of 4 bins qualifies as an interface if it contains two bins of each identity and satisfies an abundance threshold for select ligand–receptor pairs (**Methods**). Ligand-receptor pairs used here are limited to those with statistically significant colocalization tissue-wide (as measured by the communication z-score), leading to added specificity of interface prediction. Detection parameters (grid size, abundance threshold) are hand-tunable to accommodate variation in resolution and gene panel composition from one data set to another.

The above two steps achieve the interface prediction task at the core of the methodology. Also included in SCOUBI are downstream functionalities that build on the resulting interface map: 1) contrast transcriptomic profiles of interfaces to those of the broader extrasomatic background (**Figure 1e**), 2) if cell type or brain region annotations are provided, link each interface to its proximal cellular context and compare interface transcriptomes across regions and cell types (**Figure 1f)**, and 3) contrast interface expression profiles against somatic profiles, allowing systematic identification of transcripts selectively enriched at interfaces (**Figure 1g**).

### A map of axon- and dendrite-enriched loci across the mouse hemibrain

We used the SCOUBI framework to first analyze a Xenium Prime mouse hemibrain dataset (Genomics 2024) comprising 5,027 genes profiled across 63,162 cells (**Supplementary Fig. 1**). Binning the tissue at 2×2 *μ*m resolution (see below and **Methods**) yields > 6.1M spatial bins with non-zero expression, of which ∼30% overlap with segmented cell bodies and were excluded from further analysis (**Supplementary Fig. 2**). The remaining ∼4.2M bins (70%) constitute the unsegmented extrasomatic space, harboring ∼42% of all detected transcripts. These extrasomatic bins were subjected to neurite classification with SCOUBI using 13 axonal and 13 dendritic marker genes that comprise literature-reported markers of processes as well as pre- and postsynaptic compartments (**Supplementary Table 1**).

The trained model produces a tissue-wide map in which 27% and 25% of non-zero-expression extrasomatic bins are classified as axon- and dendrite-enriched, respectively. (**Figure 2a, Supplementary Fig. 2, Methods**). These labels represent local neurite-type enrichment rather than discrete anatomical assignments, since axons and dendrites are intermingled throughout neuropil. A primary visual validation of the model’s annotations comes from their concordance with expected tissue organization. The fiber tract shows axon-enriched assignments (**Figure 2b**), whereas other regions such as the cortex display a more spatially heterogeneous mix of axonal and dendritic bins (**Figure 2c**). This pattern is consistent with the known organization of white matter, which is dominated by bundled axonal fibers (Matute and Ransom 2012), and gray matter, where axonal and dendritic processes are densely packed and intermingled (Gao et al. 2019). (Also see **Supplementary Note 1, Supplementary Figure 3**.)

**Figure 2:**
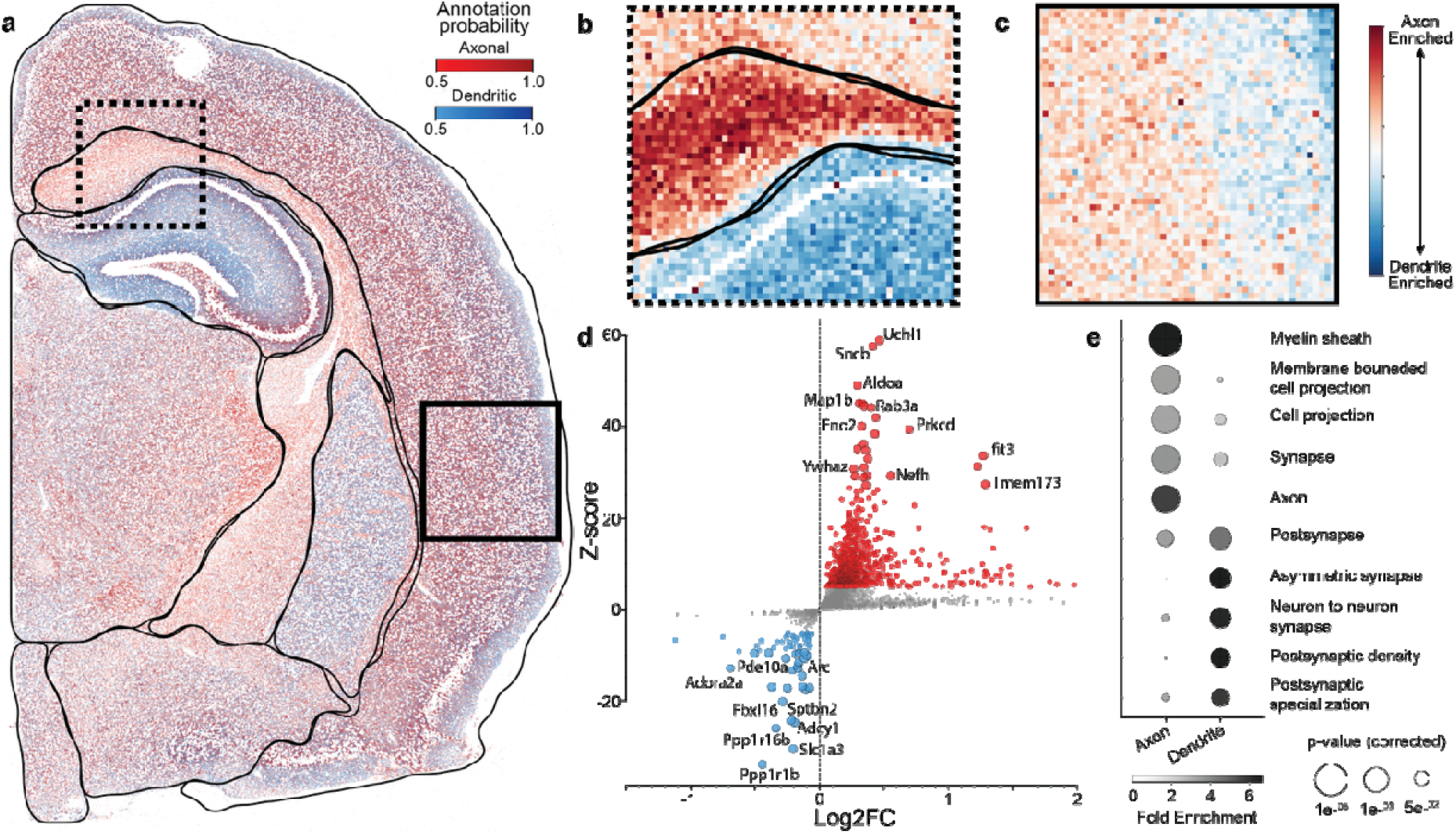
SCOUBI generates a tissue-wide neurite annotation map of the mouse hemibrain. **(a)** Tissue-wide neurite classification map of a mouse hemibrain (Xenium Prime data). Bins are colored red for axon-enriched or blue for dendrite-enriched assignments, with color intensity scaled by classification confidence. Regional boundaries are overlaid manually. **(b, c)** Zoomed views illustrating concordance between SCOUBI annotations and expected tissue organization. For visualization, bins were aggregated into µm windows and color intensity reflects the proportion of axonal and dendritic bins in each window. The fiber tract region (b) shows predominantly axon-enriched assignments, consistent with the organization of white matter as bundled axonal fibers. A cortical gray matter region (c) shows a more heterogeneous mixture of axon- and dendrite-enriched bins, consistent with the dense intermingling of axonal and dendritic processes in neuropil. **(d)** Scatter plot of log_2_ fold-change versus z-score (two-sided proportion test) quantifying differential expression of (non-marker) genes between axonal and dendritic bins. Genes with |z|>5 are colored in red (enriched in axonal) or blue (enriched in dendritic). Known axonal genes (*Uchl1, Map1b, Rab3a*) and postsynaptic genes (*Ppp1r16b, Adcy1, Sptbn2*) are enriched in their expected neurite classes. **(e)** Overrepresentation analysis of the top 25 enriched genes for each neurite class from (d), confirming enrichment for class-appropriate biological processes.

Although individual neurite diameters fall below the 2×2 *μ* m bin resolution of the map, marker gene transcripts were accurately assigned to the expected bin type. For instance, an axonal marker gene’s transcripts were found in axonal bins 83% of the time, averaged across all axonal markers. The corresponding number for dendritic markers was 90% (**Supplementary Fig. 4**). At larger bin sizes, as expected, each bin becomes increasingly heterogeneous in its neurite content, leading to a corresponding loss of accuracy (**Supplementary Note 2, Supplementary Fig. 5**). We next identified genes beyond the input marker set that are differentially distributed between annotated axonal and dendritic-enriched bins (two-sided proportion test, |z-score| > 5) (**Figure 2d, Supplementary Data 1**). Among the most significant axon enriched genes are *Uchl1, Map1b*, and *Rab3a*, each with well-characterized presynaptic or axonal functions (Meixner et al. 2000; Leenders et al. 2001; Liu et al. 2019), while dendritic-enriched genes include *Ppp1r16b, Adcy1*, and *Sptbn2*, genes with known postsynaptic roles (Magdaleno et al. 2002; Stankewich et al. 2010; Zhang and Wang 2023). Overrepresentation analysis of the top 25 most significant genes in each category revealed enrichment for class-appropriate biological processes (**Figure 2e, Supplementary Data 2,3**), providing orthogonal support for the accuracy of our bin annotations at scale.

### Interfaces mirror spatial distribution of neuronal communication

Having established axonal and dendritic bin annotations in extrasomatic space across the tissue section, we next identified interfaces by scanning for loci of axonal–dendritic convergence that express statistically shortlisted ligand–receptor pairs (**Methods**). This procedure yielded ∼32K interfaces comprising ∼68K bins (roughly 1% of non-zero-expression extrasomatic bins and 3% of axonal/dendritic labelled bins, **Supplementary Figure 2**).

We classified the identified interfaces according to the presence of excitatory or inhibitory presynaptic markers and noted that excitatory interfaces outnumbered inhibitory interfaces by approximately fourfold (**Supplementary Fig. 6a**), consistent with the established predominance of excitatory synapses in the mouse brain (Sievers et al. 2024). Interfaces were detected across all brain regions, with their relative abundance being highest in the hippocampus and lowest in fiber tract regions (**Figure 3a, Supplementary Fig. 6b**). This spatial distribution is consistent with established estimates of excitatory synapse density across the mouse brain, which place the hippocampal formation and cortex among the most synapse-dense regions, and white matter tracts among the least (Santuy et al. 2020). We then examined whether finer-grained interface distributions reflect known synaptic architecture, focusing first on the hippocampus. A contour map visualization (**Figure 3b**) revealed that interface density is highest in the CA2 and CA3 subfields and progressively lower toward CA1 and the dentate gyrus. This regional ordering corresponds to the well-established gradient of synaptic convergence across hippocampal subfields, with CA2 and CA3 representing regions of high synapse density, CA1 receiving a comparatively restricted set of inputs, and the dentate gyrus known to have the

**Figure 3:**
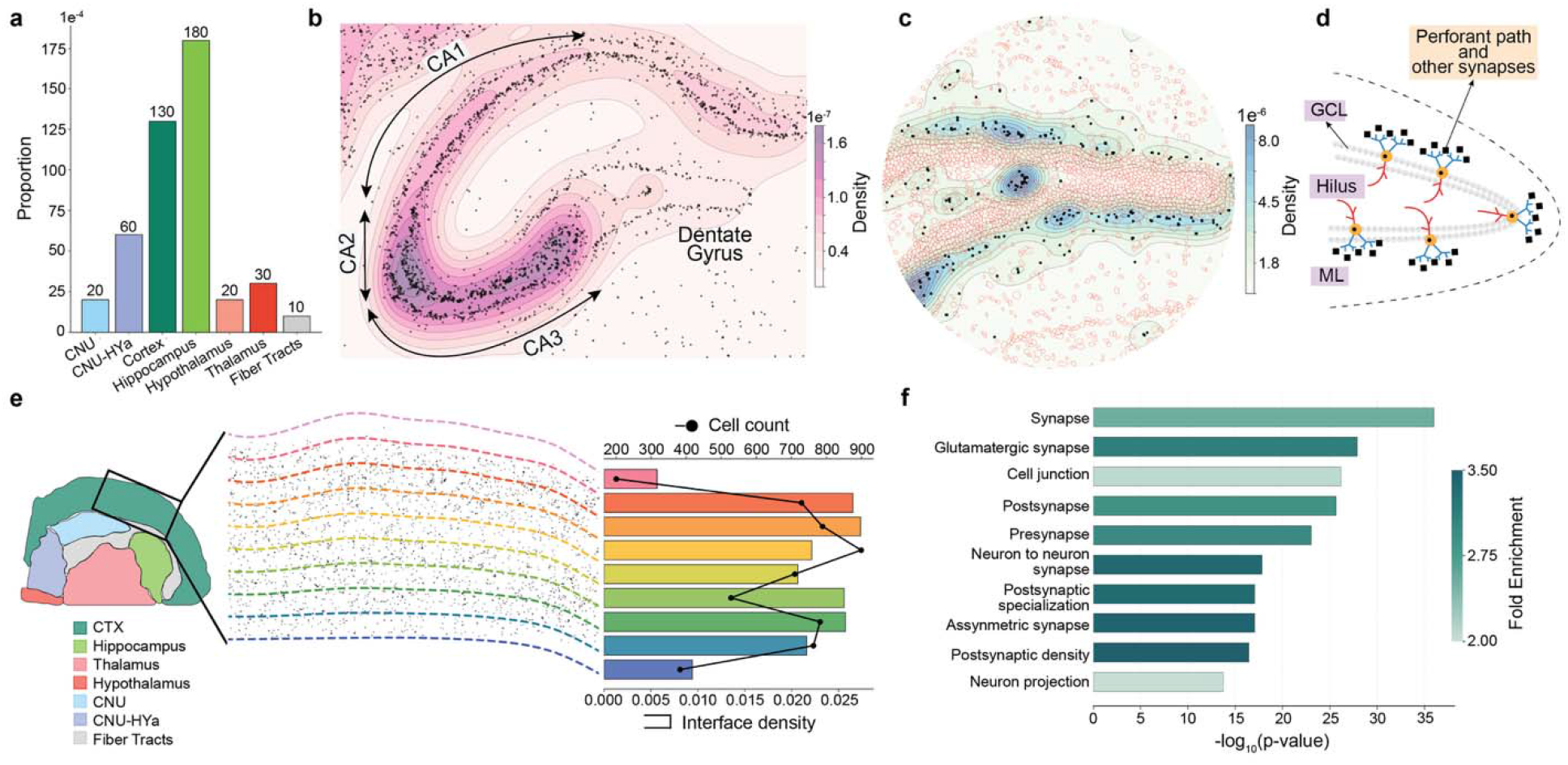
SCOUBI identifies interfaces with anatomically consistent spatial distributions and synapse-enriched transcriptomic composition. **(a)** Interface abundance (count of interface bins divided by total bins in a region) across brain regions. Abundance is highest in the hippocampus and lowest in fiber tract regions, consistent with known synaptic density estimates across the mouse brain. **(b)** Interface distribution (black dots) across hippocampal subfields, visualized with kernel density estimation (color gradient) and contour lines. Interface density is highest in CA2/CA3 and decreases toward CA1 and the dentate gyrus, mirroring the established gradient of synaptic convergence across the hippocampal formation. **(c)** Magnified view of the dentate gyrus from (b), with segmented cell bodies shown in red outline. **(d)** Schematic illustration of dentate gyrus projection architecture. Granule cell somata reside in the granule cell layer (GCL), extend axons (red) through the hilus toward CA3, and dendrites (blue) into the molecular layer, where they receive synaptic input from incoming axons. **(e)** Interface abundance and segmented cell body counts across ten successive radial segments of a representative cortical section. Interface fraction peaks around layers 5/6 and is elevated in superficial layers, closely tracking cell body distribution and consistent with known laminar synaptic organization in mouse cortex. **(f)** Analysis of 325 genes significantly overrepresented at interfaces relative to library-size-matched extrasomatic background bins (z-score > 3, p < 1×10^−4^), revealing enrichment for synapse-associated structures and molecular components.

lowest synaptic density overall (Amaral et al. 2007; Santuy et al. 2020). We also detected interfaces along the molecular layer of the dentate gyrus (**Figure 3c**) – a region known to harbor synapses between axons projecting from regions such as the entorhinal cortex (perforant path), and dendrites of granule cells (Amaral et al. 2007) (**Figure 3d**). Finally, to assess whether interfaces capture laminar synaptic organization in the cortex, we divided a representative cortical section into ten equally spaced radial segments and computed the fraction of bins assigned to interfaces within each segment (**Figure 3e**). This fraction rises through the superficial layers, dips around layer 3, peaks around layer 5, and then declines toward the white matter boundary. This pattern closely mirrors the arrangement of neuronal cell bodies across cortical segments and is consistent with the known laminar organization of cortical synapses, where synaptic puncta density is elevated in layers 2/3 and layer 5 in the mouse somatosensory cortex (Gao et al. 2019).

### Interface transcriptomes align with synaptosome profiles

We next asked whether and how interface bins differ from the broader extrasomatic space in terms of transcriptomic composition. For each gene, we computed the proportion of interface bins with detectable expression of the gene and assessed its statistical significance against a null distribution reflecting background extrasomatic bins. (The test ensures that any observed enrichment reflects compositional specificity rather than transcript density differences, see **Methods**). Genes significantly enriched at interfaces (325 genes, z-score > 3, p-value < 1×10^−4^, **Supplementary Data 4**) were strongly associated with synapse-associated structures and functions (**Figure 3f, Supplementary Data 5**), establishing that interfaces are compositionally distinct from the surrounding extrasomatic space in a manner consistent with their inferred biological identity.

To further assess the biological relevance of interface-enriched genes, we compared them to gene sets derived from prior synaptosome transcriptome and translatome profiling studies (**Supplementary Table 2**). Overlap between gene sets was quantified using a one-sided hypergeometric test (**Methods**). Interface-enriched genes show significant overlap with synaptosome-derived gene sets across multiple studies (**Figure 4a, Supplementary Data 6**). These overlaps are stronger than those exhibited by genes with the highest total expression across the extrasomatic space (“top extrasomatic”). The main exception was a gene set from Glock et al. (Glock et al. 2021), who profiled the broader neuropil translatome from microdissected neuronal processes rather than a purified synaptosome fraction, possibly explaining the comparatively strong overlap with genes broadly abundant in extrasomatic space. We also noticed that the fold enrichment between interface-enriched genes and synaptosome gene sets exceeds the overlap observed among the different synaptosome studies (**Figure 4b**). These results, along with our findings above, indicate that interfaces are enriched for genes associated with synaptic compartments, supporting the interpretation that SCOUBI isolates molecular programs linked to intercellular signaling rather than broadly distributed extrasomatic RNA.

**Figure 4:**
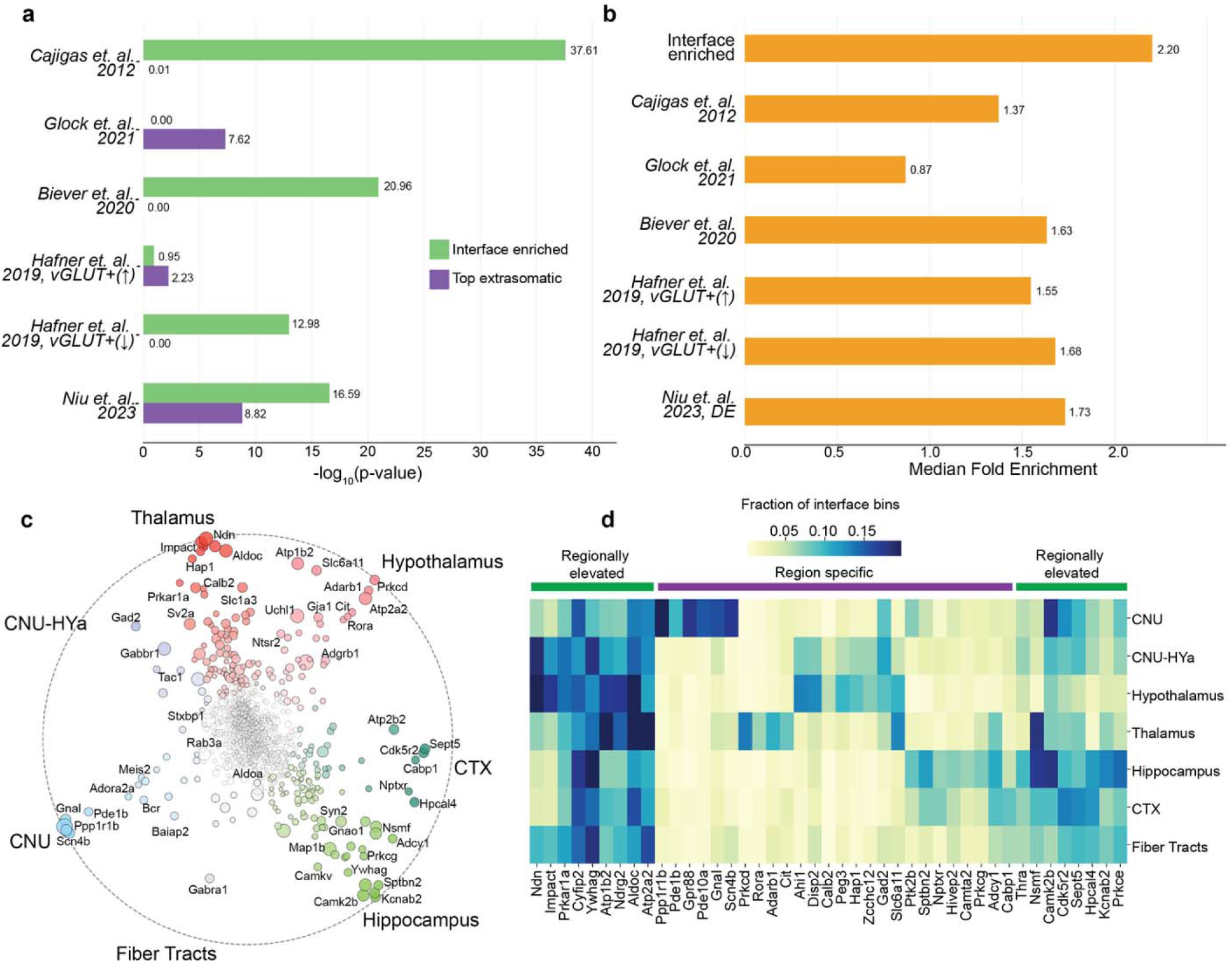
Interface transcriptomes align with synaptosome profiles and capture a conserved synaptic core elaborated by region-specific molecular signatures. **(a)** Bar chart showing the negative log p-values (Hypergeometric test) of overlap between synaptosome profiling derived gene sets (six studies) and either interface-enriched genes (green) or an equal number of top extrasomatic-expressed genes (purple). Overall, interface-enriched genes show significantly stronger overlap with synaptosome profiles than genes selected purely based on extrasomatic expression. **(b)** For each given gene set, hypergeometric fold enrichment was computed against every other gene set. Bars show the median of those pairwise enrichment values for that gene set. Interface-enriched genes show greater overlap with synaptosome-derived gene sets than synaptosome-derived gene sets show with one another. **(c)** RadViz visualization of genes, by their expression in interfaces of different brain regions. (Neurite marker genes were excluded.) Highlighted genes span key components of synaptic and intercellular communication, including vesicle release machinery (*Syn2, Sept5*), cytoskeletal and scaffolding elements (*Sptbn2, Ywhag*), calcium signaling regulators (*Camk2b, Hpcal4*), ion channel regulators (*Scn4b, Kcnab2*), and perisynaptic glial factors (*Gja1, Atp1b2*). Pan-synaptic genes (*Rab3a, Stxbp1, Aldoa*) occupy the plot center while region-displaced genes cluster toward hippocampal, thalamic, cortical, and striatal poles, exemplifying regional transcriptomic diversity superimposed on a core synaptic program. **(d)** Heatmap of region-enriched interface genes, revealing two patterns: genes broadly expressed but elevated in some regions, and genes with near-exclusive enrichment in a single region. CNU interfaces are enriched for a coherent dopamine signaling axis comprising *Gpr88, Ppp1r1b, Pde10a, Pde1b*, and *Gnal*, consistent with dopamine–glutamate signal integration at striatal synapses. Thalamic interfaces are enriched for genes with established thalamic expression including *Adarb1, Prkcd, Cit*, and *Rora*, the latter consistent with synaptic mRNA trafficking and local translation programs.

### Interface transcriptomes exhibit regional diversity

The above findings raise the question of whether interface transcriptomes exhibit region-specific molecular signatures, as opposed to a spatially uniform synaptic program regardless of anatomical context. To answer this, we computed for each gene the fraction of interface bins expressing that gene, separately for each brain region, and visualized these through RadViz (Hoffman et al. 1999) (**Figure 4c, Supplementary Data 7, Methods**). (Marker genes used for neurite classification were excluded, to avoid circularity.) A substantial fraction of examined genes exhibited a region-skewed level of interface-expression, including genes involved in synaptic and intercellular communication, spanning vesicle release machinery, scaffolding and cytoskeletal components, calcium signaling regulators, ion channel and transport genes, and perisynaptic glial factors. Known examples of pan-synaptic genes, such as *Rab3a* and *Stxbp1*, were placed near the center of the plot, meaning that no strong region-skew was observed for these genes. This analysis suggests that interfaces harbor a conserved synaptic molecular core that is elaborated upon by rich and anatomically specified transcriptomic diversity.

A closer examination reveals that the region-specific interface genes (above) can be further categorized into two broad patterns (**Figure 4d, Methods**). The first sub-category comprises genes that are detectably expressed across multiple regions but show elevated interface abundance in specific anatomical contexts, suggesting region-specific elaboration of a broadly deployed synaptic program. The second pattern comprises genes with highly restricted interface presence, concentrated almost exclusively within a single region, suggesting the existence of molecularly distinct interface identities across the brain. To characterize these distinct molecular identities, we focused on the two regions showing the strongest and most coherent regional specificities, the Cerebral Nuclei (CNU) and the thalamus. In the CNU, the most region-specific signal is defined by *Gpr88, Ppp1r1b, Pde10a, Pde1b*, and *Gnal*, a set of genes with established roles in medium spiny neuron signaling. Together, these genes are known to define a dopamine-responsive molecular axis that modulates how glutamatergic input is integrated in striatal circuits (Svenningsson et al. 2004; Nishi et al. 2008; Herve 2011; Quintana et al. 2012). On the other hand, genes with thalamus-specific interface expression are those with previously reported involvement in thalamic circuits, including *Adarb1* (Higuchi et al. 2000; Lundin et al. 2020), *Prkcd* (Zeisel et al. 2018; Colgan et al. 2023; Yao et al. 2023), and *Cit* (Furuyashiki et al. 1999). This pattern extends to *Rora*, a nuclear receptor whose main known function is the transcriptional regulation of synaptic and neurodevelopmental gene programs (Sarachana and Hu 2013; Vitalis and Mariani 2018; Ribeiro and Sherrard 2023), with known thalamus enrichment (Ino 2004; Ortiz et al. 2020). The detection of *Rora* at interfaces is intriguing given its canonical nuclear function but can be accounted for by the documented non-nuclear localization of specific isoforms (Aschrafi et al. 2006).

The above observation regarding *Rora* raises the broader question of whether localization of transcripts encoding regulatory proteins to distal sites such as synapses is a recurring feature in the brain. Consistent with this, transcripts coding for transcription factors (TFs) were frequently detected across interfaces, with 54,428 of the 68,576 interface bins containing at least one TF transcript. Moreover, among interface-enriched genes not reported as enriched in published synaptosome profiling datasets, 17 of 63 encode TFs (p = 1.7×10^−3^, Hypergeometric test). These genes organized into distinct modules, with genes within each module exhibiting similar regional abundance patterns. Furthermore, their detection was not associated with increased proximity to segmented cell bodies (**Supplementary Note 3, Supplementary Fig. 7**). Together, these findings support the view that localization of TF-encoding mRNAs to distal sites is a widespread yet under-reported phenomenon in the mammalian brain. (This may partly reflect differences in the criteria used to define interface and synaptosome enrichment.) While their functional significance remains to be established, their extensive and anatomically patterned detection suggests a broader role for synaptic-nuclear coupling, beyond the known activity-regulated TFs such as Creb (Di Nardo et al. 2007; Cox et al. 2008; Lai et al. 2008; Ji and Jaffrey 2014).

Interfaces therefore likely represent more than sites of physical axonal–dendritic proximity – their molecular composition varies systematically across brain regions and reflects regional signaling context. (Similar regional patterns are observed for the statistically significant ligand– receptor pairs used in interface identification; see **Supplementary Note 4, Supplementary Fig. 8**.)

### Interfaces reveal localized molecular programs missed in cellular expression analysis

Interfaces are an extension of the cellular molecular environment, and it is therefore expected that genes enriched (relative to extrasomatic background) at interfaces of a given region will also be preferentially expressed in the cells of that region. With this baseline expectation, we sought to identify cases where gene enrichment in interfaces of a region goes beyond what cellular expression alone would predict, potentially leading to hypotheses about preferential mRNA localization to interfaces through mechanisms such as UTR-mediated transcript anchoring (Holt and Schuman 2013; Tushev et al. 2018). To systematically identify such cases, we calculated z-scores quantifying a gene’s presence in a region relative to that in all other regions and ranked genes according to these scores. We repeated this process three times, analyzing transcriptomes of cell bodies only, interfaces only, and the extrasomatic space only (**Methods**). Contrasting a gene’s rank in the interface-only analysis against its rank in the cell body analysis for the same region allows us to assess the degree to which its interface enrichment agrees with, or departs from, the local cellular expression landscape (**Figure 5a**, also see **Supplementary Data 8**). Genes along the diagonal reflect interface enrichment closely mirroring cellular expression, while genes departing from it reflect preferential accumulation at interfaces of a region beyond what their cellular abundance in that region would predict. We additionally restrict attention to genes not enriched in the extrasomatic space of that region versus other regions, and these form the focus of the analysis below.

**Figure 5:**
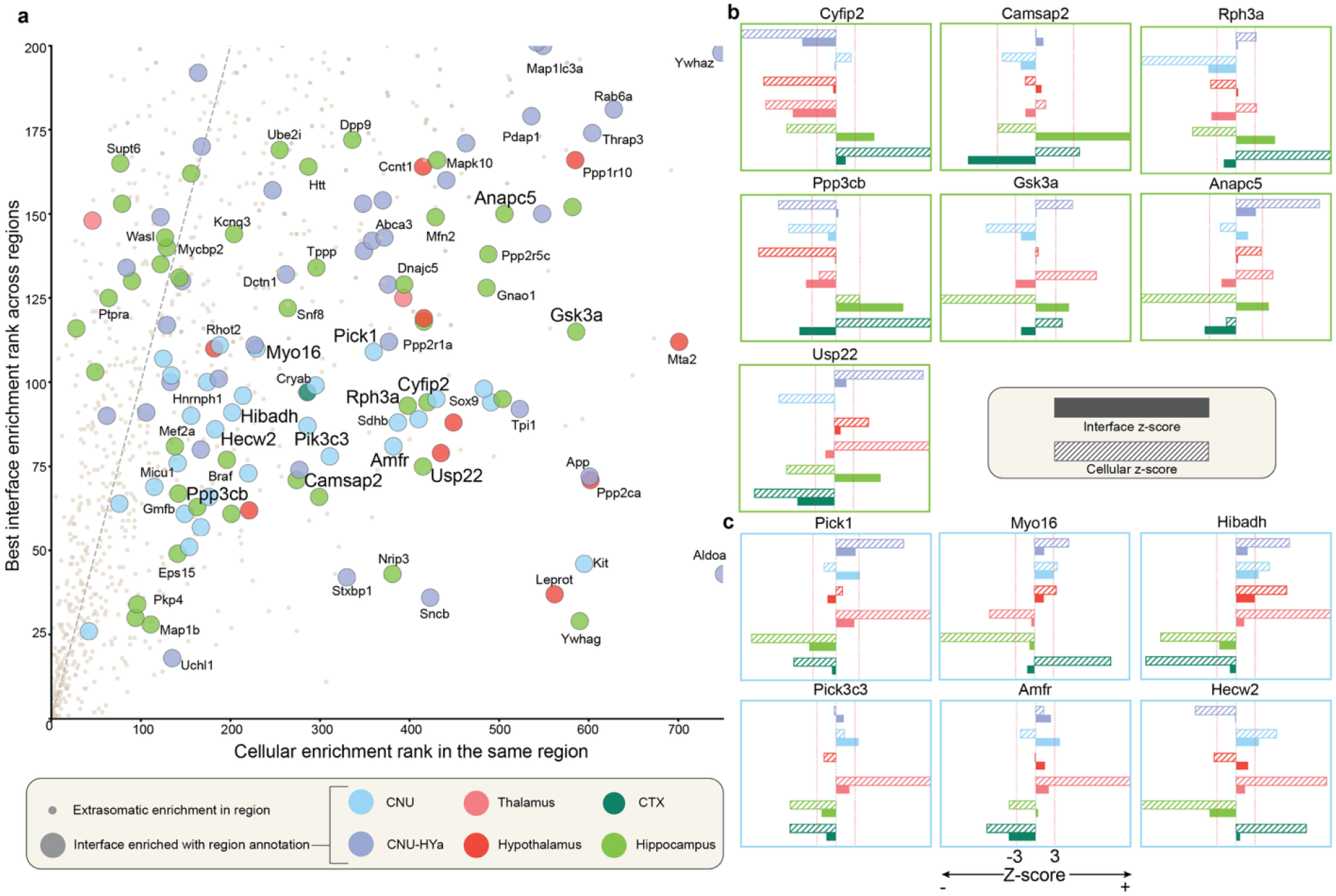
Interfaces can reveal region-specific synaptic programs not reflected in cellular expression. **(a)** Genes are scored and ranked by their enrichment in a region relative to all other regions, while restricting analysis to interfaces. For each gene, the best rank across all regions is shown on the y-axis, with color indicating the region. X-axis shows the rank of that gene in terms of its enrichment in the same region (vs. other regions), but with the analysis now being limited to cell bodies. Genes located along the diagonal reflect interface enrichment in line with cellular abundance, while placement to the right of the diagonal indicates preferential accumulation at interfaces beyond that expected from local cellular expression. Size of point/circle indicates whether the regional enrichment shown is also significant when comparing to the broader extrasomatic region. **(b, c)** Proportion test z-scores quantifying interface enrichment in the hippocampus (b) and cerebral nuclei (c). Filled bars show interface z-scores and dashed bars show cell body z-scores, with color indicating the region of enrichment. Vertical lines indicate the z-score threshold of |z| = 3 and z-scores are capped at |z| = 15. Genes shown represent cases where interface enrichment rank substantially exceeds cell body enrichment rank, pointing to preferential synaptic localization. Hippocampal genes span cytoskeletal remodeling, receptor anchoring, and synaptic elimination programs. CNU genes organize into receptor trafficking, bioenergetic support, and protein homeostasis modules.

Among hippocampal interface-enriched genes (green circles in Figure 5a) we observe a subset (**Figure 5b, Supplementary Data 9**) tied to activity-dependent synaptic remodeling, the process central to memory encoding in this region. Shown in the figure is a selection of seven genes whose interface expression is significantly enriched in hippocampus (solid light green bars), but cell body expression is mostly depleted in this region (corresponding dashed bars). These seven genes span multiple aspects of synaptic remodeling: *Cyfip2* (Lee et al. 2017; Zhang et al. 2019) and *Camsap2* (Yau et al. 2014) are associated with processes that support structural changes in dendritic spines, *Rph3a* (Stanic et al. 2015) has been implicated in the stabilization of NMDA receptors at synaptic membranes, *Ppp3cb* (Kim and Ziff 2014) and *Gsk3a* (Draffin et al. 2021) mediate long-term depression, contributing to bidirectional synaptic plasticity, *Anapc5* drives the ubiquitin-dependent elimination of weak or redundant connections (Bobo-Jimenez et al. 2017), while *Usp22* regulates the transcriptional programs that maintain ongoing plasticity (Zhang et al. 2022). Collectively, the interface enrichment of these genes highlights localized molecular programs that support synaptic remodeling and plasticity, processes central to hippocampal synaptic function.

We next examined genes enriched in interfaces of the CNU (**Figure 5c, Supplementary Data 10**), specifically, the striatum. Striatal medium spiny neurons (MSNs) are known to extend elaborate dendritic arbors densely studded with thousands of spines, creating a need for local molecular regulation far from the soma. The interface-enriched gene set identified here reflects multiple interrelated functional modules. *Pick1* is associated with the rapid sorting and turnover of surface receptors, while *Myo16* couples these synaptic changes directly to the actin-based spine structure (Nakamura et al. 2011; Roesler et al. 2019). *Hibadh* (Murin et al. 2008) and *Pik3c3* (Wang et al. 2011) are linked to metabolic support and autophagic pathways, consistent with local energetic and maintenance demands. Finally, *Amfr* (Zhang et al. 2025) and *Hecw2* (Fajner et al. 2021) are involved in protein homeostasis, contributing to the regulation of synaptic protein composition. Together, the interface enrichment of these transcripts aligns with the known demands of synaptic regulation in the striatum.

We summarize that the interface-enriched gene sets of the hippocampus and striatum are not merely reflections of the dominant cell types that populate these regions. They reveal molecularly distinct programs, potentially shaped by the specific demands each region places on its synapses, that are fundamentally inaccessible to cell-type-resolved or bulk approaches. Notably, several of these genes, including *Usp22* and *Rora* in the analyses above, have canonical nuclear functions, yet their detection in prior synaptosome profiling studies indicates that their presence at synaptic compartments is biologically grounded rather than artifactual (see **Supplementary Note 3, Supplementary Figs. 7, 9**). By resolving transcript enrichment at the level of individual interfaces across tissue, SCOUBI makes this layer of regional molecular organization directly analyzable.

### Reproducibility, benchmarking and ablation analysis of SCOUBI

We next asked whether the interface transcriptomic signatures reported above are reproducible across independent datasets. We examined genes enriched in interfaces identified by SCOUBI in six additional wild type mouse brain sections profiled using Xenium in three studies – Crowley et al. (Crowley et al. 2026), Uenishi et al. (Uenishi et al. 2026) and Forero et al. (Forero et al. 2025). These included three coronal sections from Crowley et al., male and female coronal hemispheres from Uenishi et al., and a sagittal section from Forero et al. (The studies used largely identical Xenium 5K gene panels.) For Crowley et al., gene-level enrichments were considered if observed in at least two of three replicates, while for Uenishi et al., enrichment was required in both male and female sections. Comparing interface-enriched genes from the resulting four study-level sets revealed 45 such genes shared across all studies (SuperExactTest p-value = 1.05 × 10^-150^, **Figure 6a**). (Neurite markers and ligand–receptor genes used for interface identification were excluded from this analysis.)

To assess the biological relevance of genes consistently reported as interface-enriched across studies, we compared them with SynGO (Koopmans et al. 2019) and the synaptosome-derived gene sets examined above. Among genes reported in at least two, three or all four studies, 259 of 289, 136 of 152 and 42 of 45 genes, respectively, were represented in these reference sets (**Figure 6b**). These overlaps were significantly greater than expected by chance across all thresholds (p-values < 10^−12^, **Supplementary Data 11**), indicating that the association of interface transcriptomes with synaptic biology is consistently recovered across independent datasets.

**Figure 6:**
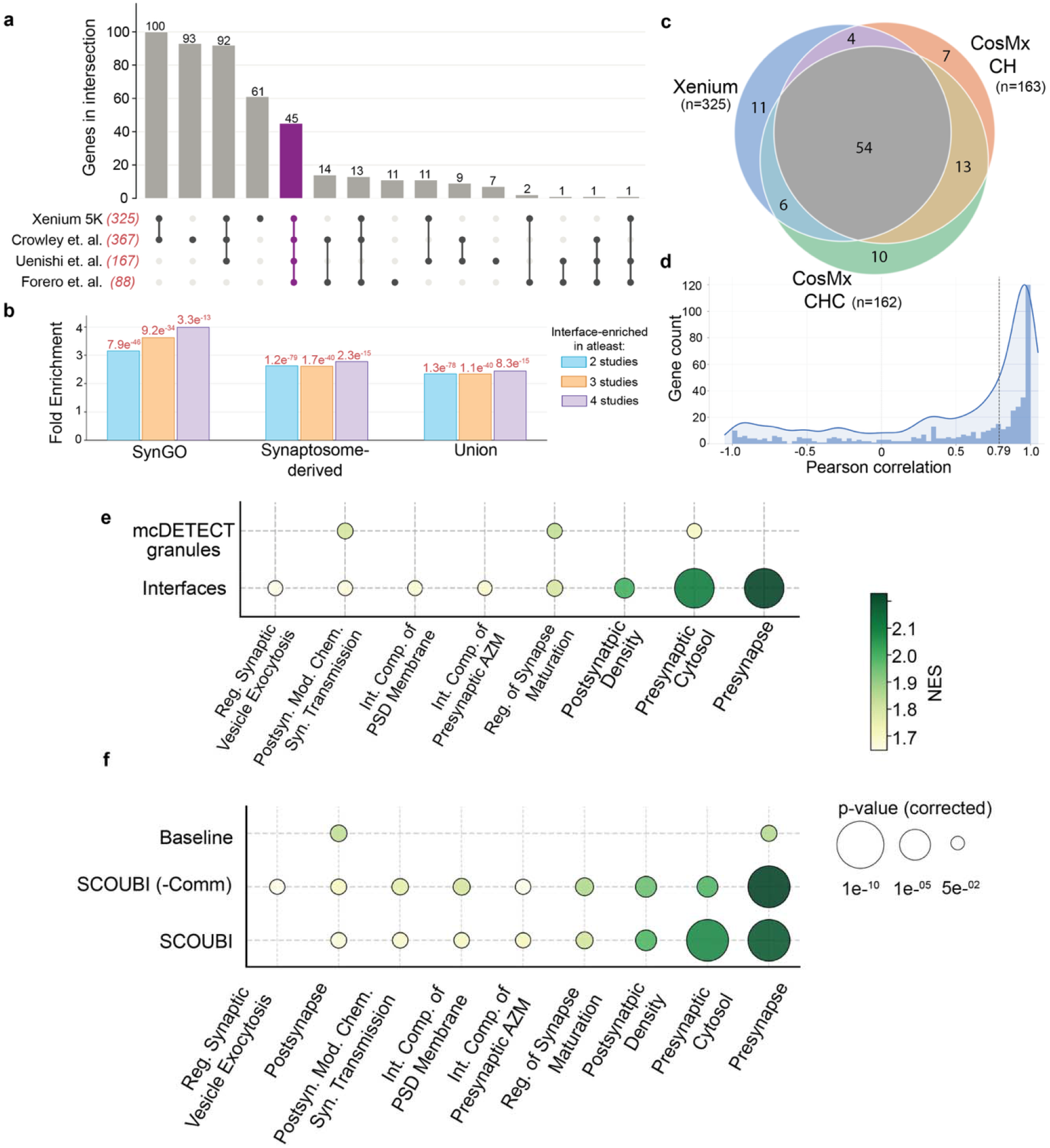
SCOUBI-predicted interfaces are reproducible across datasets and capture synaptic molecular organization more than existing approaches. **(a)** UpSet plot showing the overlap of interface-enriched genes across four independent studies comprising seven mouse brain sections profiled using Xenium. (Gene set sizes shown in red parentheses.) Only genes measured in all studies were considered, excluding neurite markers and ligand–receptor genes. The shared intersection of 45 genes is highlighted in purple. **(b)** Fold enrichment of interface-enriched genes shared across at least two, three or all four Xenium studies in SynGO, synaptosome-derived gene sets and their union. Enrichment was assessed using a one-sided hypergeometric test against the 4,941 genes measured across all sections, with corresponding p-values indicated above the bars. **(c)** Overlap of interface-enriched genes (relative to extrasomatic space) found in the Xenium and two CosMx datasets. Strong cross-dataset overlap shows that the interface transcriptomic signature is reproducible across imaging platforms and independent tissue samples. **(d)** Distribution of per-gene Pearson correlations of regional expression profiles between CosMx CH and Xenium datasets; expression values were based on detected interfaces in each of three major regions. High correlations for most genes indicate that anatomical patterning of interface transcriptomes is a stable tissue property rather than a platform or sample-specific artifact. **(e)** Gene set enrichment analysis of genes enriched in interfaces or in mcDETECT granules relative to extrasomatic background. Interfaces show substantially stronger enrichment for synapse-related terms than granules do, reflecting the specificity gained by requiring axonal-dendritic convergence over transcript aggregation alone. **(f)** Gene set enrichment analysis of genes enriched at interfaces relative to the extrasomatic background using three approaches: the full joint “SCOUBI” objective, SCOUBI without the communication loss “SCOUBI (-Comm)”, and a “Baseline” that assigns neurite identity from average axonal and dendritic marker expression. Interface-enriched genes derived from the model trained using the joint objective show the strongest enrichment for synapse-related terms. In both (e) and (f), color indicates normalized enrichment score (NES), and only enriched terms with corrected p-value < 0.05 in any of the methods are shown.

We next investigated whether these findings are preserved across imaging technologies. We applied the SCOUBI method to two independent CosMx datasets of the mouse brain, a coronal

hemisphere (CH) section and a coronal hippocampus and cortex (CHC) section. Both datasets were preprocessed identically to the Xenium data (above) and binned at 2×2 *μ* m resolution. The CosMx gene panel comprises 950 genes, of which 558 are shared with the Xenium dataset. Six of the 13 axonal markers and 7 of the 13 dendritic markers used for neurite classification above are represented in the CosMx panel (**Supplementary Table 1**). Consistent with the reduced axonal marker representation, a lower fraction of bins is called axonal relative to the Xenium data. The fraction of dendritic bins decreases in the CH dataset but increases in the CHC, likely reflecting the dendritic organization of hippocampal subfields emphasized in the latter. Consistent with this, interface abundance is higher in the CHC data, where the limited coverage of fiber tract regions raises the overall interface fraction (**Supplementary Figs. 10,11**). Despite these differences in panel composition and section anatomy, axonal and dendritic annotations in the CosMx data sets recapitulate the neuroanatomical organization observed in the Xenium hemibrain. The dentate gyrus and thalamus display the same morphological patterning, indicating that neurite classification using SCOUBI is robust across platforms and gene panels (**Supplementary Fig. 12**).

We next asked whether interfaces in the CosMx datasets are molecularly distinct from the broader extrasomatic space in the same way as reported above. Repeating the statistical procedure from above, we obtained 163 and 162 interface-enriched genes in the CH and CHC datasets respectively (z-score > 3, p < 1×10^−4^, excluding marker and ligand-receptor genes) (**Supplementary Data 12,13**), with gene set analysis again recovering synaptic terms (**Supplementary Data 14,15**). Of these, 136 genes are shared between the two CosMx datasets (hypergeometric p = 9×10^−104^), confirming that the interface transcriptomic signature is reproducible across independent CosMx samples. Comparing these genes to those enriched at interfaces in the Xenium data reveals strong overlap with both CosMx datasets (CHC: hypergeometric p = 2.9×10^−43^; CH: hypergeometric p = 2.2×10^−42^; **Figure 6c**), establishing that the molecular characteristics of predicted interfaces are reproducible across imaging platforms and studies.

Having confirmed that interfaces remain compositionally distinct from extrasomatic background, we next determined whether their region-specific transcriptomic organization is preserved across datasets. For each gene, we computed an “abundance vector” capturing its interface expression across brain regions and compared these vectors between the CosMx and Xenium datasets using Pearson correlation (**Methods**). Even across imaging technologies and independent tissue samples, regional interface expression profiles are strongly correlated for most genes (**Figure 6d, Supplementary Fig. 13, Supplementary Data 16**). The CosMx datasets identified many of the same interface-associated genes observed in the Xenium analysis, including *Ywhag* and *Aldoa*, as well as region-specific signatures such as *Rora* in the thalamus and *Ppp1r1b* and *Gnal* in the CNU (**Supplementary Figs. 14,15, Supplementary Data 17,18**). These results demonstrate that the axonal and dendritic annotations, the molecular distinctiveness of interfaces, and their anatomically constrained transcriptomic diversity are consistent features of ST data that reflect genuine biological organization rather than technical or sample-specific variation.

We next compared the SCOUBI method to mcDETECT (Yuan et al. 2025), a method that uses density-based clustering of synaptic marker genes to identify RNA “granules” in extrasomatic and distal compartments. While the goals of mcDETECT are different from ours, we included this comparison as it is the only other tool today tailored to global analysis of extrasomatic transcripts in ST data. Applying mcDETECT to the Xenium dataset using the same neurite marker set as above, we identified ∼130K granules, of which ∼56K overlapped with segmented cell bodies and were excluded from subsequent analysis. We then identified genes enriched in granules relative to the extrasomatic space using the same statistical framework as above. Granules yield 147 enriched genes compared to 325 at interfaces, and gene set analysis revealed substantially stronger enrichment of synapse-related terms in interface-enriched genes than in granule-enriched genes (**Figure 6e**). Together, these comparisons indicate that requiring axon–dendrite convergence and ligand–receptor colocalization yields loci that are more specifically enriched for synaptic programs than density-based transcript aggregates. (See **Supplementary Note 5, Supplementary Fig. 16** for a complementary analysis using troutpy (Salas et al. 2025).)

To assess the key methodological choices of SCOUBI, we compared its results with two ablated versions: one where the marker weights used in neurite classification are optimized without jointly examining communication patterns, and one where the marker weights are not optimized, i.e., axon/dendrite annotation is based on average marker gene expression (with suitable normalization) of respective class. Our joint optimization objective produced more biologically coherent neurite annotations than either ablation, with stronger neurite-specific functional enrichment, higher ligand–receptor communication scores, and improved recovery of synapse-related programs at inferred interfaces. (**Figure 6f, Supplementary Note 6** and **Supplementary Fig. 17**). These results establish the advantage of the full SCOUBI objective and show that incorporating communication-informed spatial constraints improves neurite annotation beyond optimization of marker-gene expression alone.

## Discussion

The extrasomatic fraction of imaging-based spatial transcriptomics (iST) data has often been treated as a source of noise arising from imperfect cell segmentation and ambient transcript diffusion (Salas et al. 2025), and thus discarded. Our work reveals the biological value of this substantial but largely ignored data. By imposing prior knowledge of neuronal signaling mechanisms onto this unassigned space, we show that extrasomatic transcript distributions manifest a layer of subcellular organization that is recoverable at tissue scale without specialized dissection or biochemical fractionation. The recovered “interfaces” recapitulate established neuroanatomical organization, are compositionally distinct from the surrounding extrasomatic space, and carry region-specific transcriptomic signatures that are reproducible across imaging platforms and independent tissue samples.

A key methodological innovation is how the two scoring axes – neurite identity (axonal vs dendritic enrichment) and ligand-receptor enrichment – inform and depend on each other. Neurite identity can in principle be inferred from marker gene expression in the extrasomatic space, but without supervision, relative weights of different markers are unknown. SCOUBI solves this problem by exploiting the spatial co-localization of ligand-receptor (L-R) pairs at neurites. At the same time, it is not clear *a priori* which L-R pairs should be used for this purpose, but if reliable neurite annotations are available, L-R pairs can be statistically shortlisted by exploiting co-localization with neurites. In short, neurite classification requires knowing relevant L-R pairs, which in turn requires accurate neurite annotation. Rather than treating this circularity as a limitation, SCOUBI resolves it through joint optimization in which marker-weighted neurite labeling and L-R communication scoring iteratively inform one another until convergence.

Unlike synaptosome profiling, which captures a bulk transcriptional snapshot from dissociated tissue, interfaces preserve spatial context – where signaling occurs and which cell populations are proximal – enabling inquiries about regional and anatomical specificity that are difficult to address with fractionation-based approaches. Moreover, our analytical framework adds an orthogonal dimension to current spatial transcriptomics analysis, which has largely focused on improving cell segmentation, identifying spatially variable genes, and inferring cell-cell communication (Liu et al. 2026). By positioning the extrasomatic space as a meaningful target of analysis, the SCOUBI framework exposes a layer of molecular organization that is not captured by cell-centric approaches. This is especially consequential in neural tissue, where the rules governing which mRNAs are transported into neurites, how far they travel, and under what conditions they are translated remain poorly understood. The identification of transcripts that accumulate at interfaces beyond what local somatic expression predicts suggests active and selective mechanisms of transport or local stabilization (Fernandopulle et al. 2021), connecting to a large body of work on RNA localization signals, UTR-mediated targeting, and RNA-binding protein specificity (Holt and Schuman 2013; Tushev et al. 2018; Mendonsa et al. 2023). SCOUBI generates testable hypotheses about which transcripts may be subject to such regulation and in which anatomical contexts, directly from the transcriptomic data.

Several limitations of the current framework should be noted. Extrasomatic transcript distributions capture a composite signal arising from active transport, local translation, passive diffusion, and platform-specific technical noise, including off-target probe binding, which are not disentangled here. Empirical null modeling and cross-platform validation limit false positives but resolving the individual contributions of diverse biological mechanisms remains an area for future work. Statistical performance is influenced by transcript density and gene panel composition, as highlighted by reduced marker representation in the CosMx datasets, and application to sequencing-based spatial transcriptomics remains restricted by lower RNA capture efficiency (Ge et al. 2025). A related consideration concerns how interfaces differ from transcript “granules” (Yuan et al. 2025). Though both exploit neurite marker distributions, granules are identified from local transcript clustering alone, whereas interfaces incorporate a communication score requiring the co-occurrence of axonal and dendritic signals alongside ligand-receptor enrichment, necessitating a broader spatial window. Interfaces are therefore best understood not as structural synapses but as a pragmatic approach to probing synaptic biology in iST data.

The most immediate opportunity lies in applying the framework to disease contexts where synaptic dysfunction is a central pathological feature. Neurodegenerative and neurodevelopmental disorders are associated with region-specific alterations in synaptic composition and local translation (Fernandopulle et al. 2021; Gamarra et al. 2021) – features that interfaces are positioned to capture. Realizing this potential will require gene panels that extend beyond cell type markers to include transcripts implicated in synaptic pathology, a design consideration that current imaging panels, optimized primarily for cell type identification, do not prioritize. As spatial transcriptomics technologies continue to expand gene coverage and improve sensitivity (Cohen et al. 2025), larger panels will enable more comprehensive ligand-receptor communication modeling and finer dissection of interface molecular programs across brain regions and disease states. In this context, the extrasomatic layer is likely to become increasingly useful for studying subcellular organization and disease-associated changes in local signaling.

## Methods

### SCOUBI algorithm

Identifying putative synaptic interfaces proceeds in two stages. The first stage classifies spatial bins as either axon- or dendrite-enriched (in short, “axonal” or “dendritic”) based on the expression of curated marker gene sets. The second stage identifies loci of axon–dendrite convergence enriched for putative signaling activity. Both stages operate on binned spatial transcriptomics (ST) data, where each bin *i* ∈ {1,…, *N*} is represented by its transcriptomic profile. By default, the tissue is divided into 2×2 *μ*m bins, a size chosen to minimize bin area while still capturing sufficient signal and preserving marker gene specificity (see also **Supplementary Note 2** for further details of this choice).

### Neurite classification

Let *G*^*(a)*^ and *G*^*(d)*^ denote the axonal and dendritic marker gene sets, containing *m*_*a*_ and *m*_*d*_ genes, respectively. SCOUBI employs a lightweight two-branch linear classifier. Each branch receives its respective marker gene expression vector 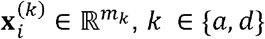, and projects it into a non-negative scalar logit via a bias-free linear layer followed by a ReLU activation and dimensionality normalization:

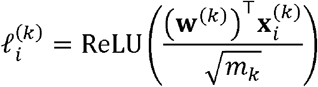

where 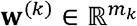 is the learned weight vector for class *k*. The two logits are concatenated to form 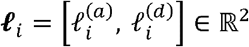 and passed through a temperature-scaled softmax:

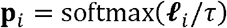

where 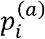 and 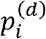 are interpreted as the probability of bin being axonal or dendritic, respectively.

The model is optimized using a combination of two loss functions, marker gene maximization loss ℒ_*gex*_ and communication loss ℒ _*comm*_. Marker gene maximization loss ℒ_*gex*_ encourages bins assigned high probability to a given neurite class to express the corresponding marker genes more strongly than the remaining bins. For each class *k* ∈ {*a*, *d*}, we compute a soft probability-weighted two-sample *t*-statistic across all bins, treating 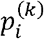 as a soft membership weight. For this, we define the in-group mean as

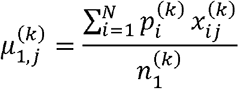

where subscript 1 indicates “in-group” (members of class *k*) and 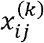 denotes the expression of marker gene *j* ∈ {1, …, *m*_*k*_} for class *k* in bin *i*. The corresponding group size, respecting the soft membership weights, is

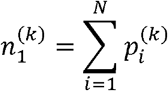

Similarly, the complementary group, i.e., non-members of class *k* (with membership weights 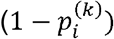, has a weighted mean and effective class size given by (respectively):

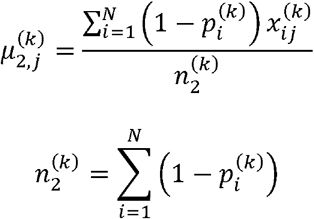

The in-group and complementary group variances are given by (respectively):

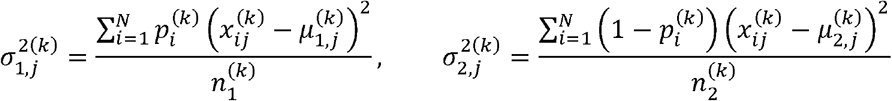

The per-gene *t*-statistic for class *k*, where *j* ∈ {1, …, *m*_*k*_} indexes individual marker genes is:

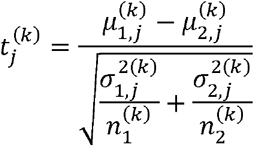

The loss term for class *k* is the negative mean *t*-statistic across its *m*_*k*_ marker genes:

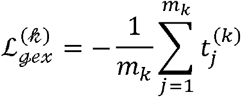

The total marker gene loss aggregates over both neurite classes:

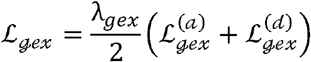

where λ_*gex*_ is a scalar hyperparameter weighting this loss term.

The communication loss ℒ _*comm*_ is an adaptation of the analytical z-score used in CellWHISPER (Kumar et al. 2026). CellWHISPER quantifies cell-cell communication strength by counting spatially proximal cell pairs in which a ligand *l* is expressed by a source cell type and its receptor *r* is expressed by a neighboring target cell type. SCOUBI applies the same principle by treating spatially adjacent axonal and dendritic bins as the source and target respectively. For each ligand–receptor pair (*l,r*)*∈P* drawn from a curated set sourced from NeuronChat (Zhao et al. 2023), we count adjacent axon–dendrite bin pairs in which the axonal bin expresses *l* and the dendritic bin expresses *r*. The observed count is evaluated against a Binomial null model that accounts for the number of eligible axon−dendrite bin pairs and the marginal expression frequencies of ligand *l* in axonal bins and receptor *r* in dendritic bins. Specifically, we compute a *z*-score *Z*_*l,r*_ quantifying the enrichment of ligand *l* and receptor *r* in neighboring axonal and dendritic bins relative to a Bernoulli null model, assuming independence among edges. This term encourages assignments where spatially proximal axonal–dendritic bin pairs are enriched for known L–R co-expression beyond random expectation. The loss is then defined as:

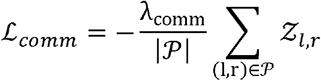

where *λ*_*comm*_ is a scalar hyperparameter. The z-score calculation is as in CellWHISPER, but the latter computes these z-scores using discrete cell-type labels. SCOUBI instead uses axonal and dendritic assignment probabilities, allowing bins and neighboring pairs to contribute fractionally according to their inferred identities. This yields a differentiable communication objective that can be optimized jointly with*ℒ*_*gex*_. We use a softmax temperature *τ* of 10^−3^ during training to obtain near one-hot assignments while preserving differentiability. (Probabilities are recomputed at a temperature of10^−1^ for visualization). An L-R pair is considered significant if its *z*-score is greater than 3 and the number of communicating edges is greater than 30, following CellWHISPER. (These criteria were met for 66 out of the 361 LR pairs tested in this work.)

The total loss is then *ℒ= ℒ*_*gex*_ +*ℒ*_*ceomm*_, which is minimized jointly to produce the final bin annotations.

### Interface identification

Interfaces are identified as 2×2 bin windows enriched for both axonal and dendritic bins and ligand-receptor expression. Two conditions have to be met: (1) 2 bins of each neurite identity must be present, and (2) at least one significant L-R pair (as defined above) must be “abundant” in the window, where an L-R pair (*l, r*) is said to be “abundant” if at least 2 transcripts of gene *l* and at least 1 transcript of gene *r* are present in the window, or vice versa.

The abundance threshold was selected based on statistical analysis using a binomial null model. Given transcript sparsity at the chosen spatial resolution, the criterion was based on bin-level detection (whether a bin has at least one transcript of the gene or not) rather than continuous abundance. For each L–R pair, (*l, r*) the global detection rate was estimated as the fraction of bins in which it is detected, denoted by *p*_*l*_ and *p*_*r*_ respectively. Under a Bernoulli null model of spatially independent detection across bins and genes,

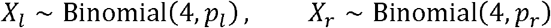

Where *X*_*l*_ and *X*_*r*_ denote the numbers of bins within a 2×2 window in which genes *l* and *r* are detected. The probability that a window satisfies the abundance criterion is,

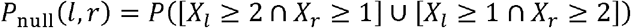

Across all three datasets analyzed, the maximum null probability *P*_null_(*l,r*) across all L–R pairs was 9.15×10^−5^ (Xenium), 2.90× 10^−5^ (CosMx CH), and 5.90×10^−4^ (CosMx CHC), confirming that the threshold is highly conservative given the observed expression sparsity.

This statistical analysis is provided as part of the SCOUBI output to guide adjustment of the threshold for datasets with different gene panel compositions or expression sparsity.

### CellWHISPER analytical z-score

SCOUBI computes ligand–receptor communication z-scores using the analytical framework implemented in CellWHISPER, adapted from cell-level communication graphs to spatial bin neighborhoods. In place of the k-nearest-neighbor graph constructed over cells in CellWHISPER, spatial adjacency was defined on the regular 2 × 2 *μ*m bin lattice, with each bin connected to its eight immediately adjacent neighbors.

Given the sparsity of bin-level transcript detection, gene expression was binarized based on the presence of at least one transcript per bin. At each optimization epoch, the model generated axonal and dendritic assignment probabilities for each bin using a low temperature softmax. This produced near-discrete neurite assignments while preserving differentiability during model optimization. For each ligand–receptor pair (*l,r*), a whisper network was constructed from adjacent axon–dendrite bin pairs. A whisper edge was defined when an axonal bin containing ligand *l* was adjacent to a dendritic bin containing receptor *r*. The observed number of whisper edges, *N*_*l,r*_ quantifies the evidence of communication for each ligand–receptor pair (*l, r*).

Communication significance was assessed using an analytical null model that preserves the marginal detection frequencies of the ligand and receptor while assuming random spatial association between their detections. The probability of detecting ligand *l* in an axonal bin was estimated as

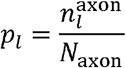

where 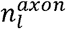 is the number of axonal bins containing ligand *l*, and *N*_*axon*_is the total number of axonal bins. Similarly, the probability of detecting receptor *r* in a dendritic bin was estimated as

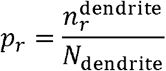

where 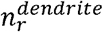 is the number of dendritic bins containing receptor *r*, and *N*_*dendrite*_ is the total number of dendritic bins. Because neurite assignments were updated during model optimization, these detection probabilities were recalculated at each epoch.

Under the null hypothesis, the probability that a neighboring axon–dendrite bin pair forms a whisper edge for ligand–receptor pair (*l,r*) is

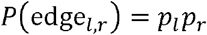

Let *N*_*E*_ denote the total number of adjacent axon–dendrite bin pairs. Under the independent-edge approximation, the number of whisper edges expected under the null model, 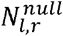, follows a binomial distribution:

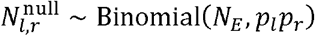

The expected null whisper edge count is therefore

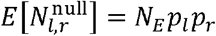

with variance

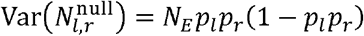

The communication z-score was calculated by comparing the observed whisper edge count with the corresponding null expectation:

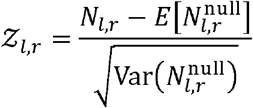

Adjacent whisper edges are not strictly independent because multiple edges may share a common bin. SCOUBI nevertheless uses the independent-edge approximation to permit efficient recalculation of communication scores throughout model optimization. CellWHISPER similarly models edge formation from the marginal expression probabilities of the two signaling genes and derives an analytical null distribution for the resulting whisper edge count.

### Enrichment analysis included in SCOUBI output

#### Empirical null construction

To characterize the transcriptome of interface bins, we compared their gene expression against an empirical background distribution drawn from extrasomatic region (ESR) bins. ESR bins are defined as those that do not overlap a segmented cell. We considered the population of ESR bins that do not belong to any interface and have non-zero total expression. From this pool, we sampled as many bins as there are interface bins (say *n*), with a library-size (total transcript count) distribution matched to that of the interface bins. The size matching is performed by extracting decile thresholds of the library-size distribution of interface bins, then sampling *n*/10ESR bins (from the above pool) in each decile range. Size-matching was undertaken so that statistical enrichment of a gene in interfaces relative to the ESR background reflects compositional specificity rather than transcript density differences between interfaces and ESR regions. The entire procedure was repeated *B*=10,000 times; each of these iterations yields a sample of the background distribution (*n* ESR bins)

#### Interface-enriched genes

For each gene, enrichment is quantified by a one-sided empirical *p*-value:

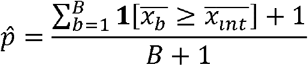

where 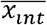 is the mean expression of the gene in the 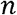 interface bins, 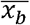 is its mean expression in the *b*^*th*^background sample of library-size-matched ESR bins,**1**[·] is the indicator function. A *z*-score is additionally computed as 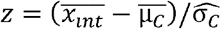, where 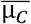 and 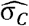 are the mean and standard deviation of 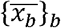. Neurite marker genes and ligand-receptor genes were excluded to avoid circularity, since interfaces are defined using these markers and their enrichment is thus expected.

### Regional enrichment comparison to cells and extrasomatic space

To identify genes enriched at interfaces within specific anatomical regions, we performed a region-wise two-sided proportions *z*-test comparing the fraction of expressing units in each region against all other regions combined. This was done independently for interface bins, ESR bins, and cells. Given the sparsity of bin-level transcript detection, expression in interface and ESR bins was binarized based on the presence of at least one transcript. Cell-level expression was binarized at the 75th percentile.

For each region *r* and gene *g*, the *z*-statistic is:

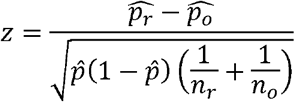

where 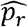 and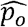 are the proportions of expressing units in region *r* and all other regions respectively, *n*_*r*_ and *n*_*o*_ are the corresponding counts, and 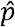is the pooled proportion.

Cell types were coarse-grained into neuronal and non-neuronal populations based on cell type labels. *z*-scores were computed separately using all cells, neuronal cells, and non-neuronal cells. Downstream analysis uses the neuronal *z*-score, as we expect the interface transcriptome to be predominantly driven by neuronal contributions.

### RadViz visualization

To characterize the transcriptional composition of interface bins across anatomical regions, we computed, for each region, the fraction of that region’s interface bins expressing each gene, yielding a gene x region expression profile matrix. Each row of this matrix constitutes the transcriptomic profile of a gene across regions. Each interface bin was assigned the regional annotation of its nearest cell.

The profile matrix is visualized using a modified RadViz (Hoffman et al. 1999) projection where regions serve as anchors on the unit circle and each gene is projected to a position determined by a softmax-weighted average of anchor positions, with the regional expression values serving as weights. Genes with a radial distance greater than 0.75 were shortlisted as having strong regional preference, which are then displayed in a hierarchically clustered heatmap for detailed visualization.

### Comparisons (performed outside of SCOUBI)

#### Comparison of interface-enriched genes with synaptosome-derived datasets

The synaptosome-derived datasets and gene-selection criteria are summarized in **Supplementary Table 2**. Transcription factors were annotated using the AnimalTFDB (Shen et al. 2023) database.

#### Comparison between independent studies

To assess reproducibility, we analyzed seven Xenium sections, six coronal and one sagittal (Forero et al. (Forero et al. 2025)). For Crowley et al. (Crowley et al. 2026), three replicates were labeled A, B and C, with enrichment required in at least two. For Uenishi et al. (Uenish al. 2026), male and female sections were labeled M and F, with enrichment required in both. The studies used largely identical Xenium 5K gene panels, with Forero et al. including an additional 100 custom genes, resulting in 4,941 genes shared across all seven sections. Only genes measured across all sections were considered, and overlap among the four resulting interface-enriched gene sets was assessed using SuperExactTest (Wang et al. 2015). To evaluate their association with synaptic reference genes, we considered genes enriched in at least two, three or all four studies. Each set was compared with SynGO, the combined synaptosome-derived reference genes, and the union of these references using one-sided hypergeometric tests. The background comprised the 4,941 genes measured across all sections, with reference gene sets restricted to this universe.

### Comparison between technologies

To assess whether region-specific interface expression was preserved across technologies, we compared Xenium with each CosMx dataset separately. For each gene detected in both datasets, we compared the fraction of interface bins expressing that gene across regions. Regions with low interface abundance were excluded, retaining the thalamus, cortex and hippocampus. Pearson correlation was then computed between the CosMx and Xenium profiles to assess cross-dataset reproducibility of region-specific interface expression.

### Comparison with other methods

mcDETECT identifies granules as spatially clustered transcripts of synaptic marker genes. However, for a controlled comparison, we ran mcDETECT (v2.0.3) (Yuan et al. 2025) on the Xenium dataset using the same neurite marker gene set (13 axonal markers and 13 dendritic markers) as that used for SCOUBI (**Supplementary Table 1**) and default parameters. Detected granules overlapping with segmented cell bodies were excluded. Although granules are reported by mcDETECT as 3D spheres, we treated them as belonging to the corresponding 2 ×2µm bin, to allow for direct comparisons.

We also compared granule bin expression against the extrasomatic empirical null using the same bootstrap framework described above. For this comparison, each granule was mapped to the 2×2µm bin containing its center, resulting in 72,606 bins for the 73,977 extrasomatic granules detected. The per-gene *z*-scores were subjected to GSEAPreranked analysis using GSEApy (Fang et al. 2023), run against the SynGO gene set and terms with FDR < 0.05 were retained.

### Preprocessing and Computational Performance

Genes with a quality score (qv) below 20 are excluded prior to model input, when quality scores are available. After binning the data, only bins in which the majority of transcripts are not assigned to any cell are retained for further analysis. Cell typing was performed via label transfer from an annotated single-cell RNA-seq reference dataset as described in Zhang et al. (Zhang et al. 2023), using the ALLCools Python package. Region annotation was performed using SPACEL (Xu et al. 2023), followed by manual correction and annotation of anatomical domains.

SCOUBI was trained on an NVIDIA H200 GPU. The Xenium dataset comprising 63,162 cells and 5,027 genes was binned at 2×2 μ*m* resolution to generate a 3,626×2,538×5,027 spatial expression tensor. Neurite classification and interface identification were completed in 11.3 minutes with a peak GPU memory usage of 49.06 GB and peak RAM usage of 48.80 GB.

## Data availability

The Xenium Prime mouse hemibrain dataset is available at https://www.10xgenomics.com/datasets/xenium-prime-fresh-frozen-mouse-brain. Additional Xenium datasets analyzed in this study are available from Crowley et al. through Zenodo (https://doi.org/10.5281/zenodo.18063582) and from Uenishi et al. and Forero et al. through GEO (GSE330686 and GSE326496, respectively). The CosMx datasets of the mouse brain coronal hemisphere (CH) section and coronal hippocampus and cortex (CHC) are available at https://brukerspatialbiology.com/products/cosmx-spatial-molecular-imager/ffpe-dataset/cosmx-smi-mouse-brain-ffpe-dataset/.

## Code availability

SCOUBI is open source and available at https://github.com/bhavaygg/SCOUBI.

## Acknowledgements

This research utilized resources from PACE at the Georgia Institute of Technology, Atlanta, USA (RRID:SCR_027619). Funding: This work was supported by the National Institutes of Health (R35GM131819 to S.S.), and Georgia Institute of Technology (Wallace H. Coulter Distinguished Faculty Chair: S.S.)

## Author Contributions

B.A. and S.S. jointly designed the study. B.A. developed the computational methods and performed the analyses. S.S. supervised the project. S.J. provided guidance and advice throughout the project. B.A. and S.S. wrote the manuscript, with input from S.J.

